# Assessment of genetic diversity patterns of different serotypes of dengue virus, prevalent in patients from Kerala, India: a strain specific mutation study and its relevance to pathogenesis

**DOI:** 10.1101/2023.02.09.527586

**Authors:** Rituraj Niranjan, T Sankari, H Manoj, A. K. Jayashree, Rameela Sanya, Basil Varghese, N. Pradeep kumar, S Muthukumaravel, Ashwani Kumar

## Abstract

The mutations are the key mechanisms responsible for virus survival and its fitness in the host. This process of mutation is implicated in the development of pathogenesis of the dengue viral disease. We report that, all the four serotypes were found to be circulating in Kerala sate of India. Serotypes 1 and 3 were the predominant type (33.3 %) compared to others. The prevalence of co-infection of serotype 1 and 4 was the highest percentage (11.1 %) compared to serotype 2 and serotype 3 (5.5 %). We further highlighted the immunopathological mechanisms of antibody neutralization, CD4^+^ T cell response and antibody dependent enhancements (ADE) for the possible cause of disease severity in coinfections. Serotype-1 does not show much variations from the previously reported strains from various regions of India. However, serotype-2 showed variations in the sequences from the other strains of serotype-2 previously reported from various regions of India and formed a distinct clade in the genotype-4. Serotype-3 and serotype-4 showed similarity with previously reported strains from India. Moreover, serotype-1 was grouping in genotype-5. Importantly, the serotype-2 grouped with genoptype-4 but exist separately. Serotype-3 was found to be grouped with the genotype-3. The serotype-4 show the very much similarities from the genotype-1 and shows little difference from the previously reported strains from India. Further, mutation in DENV-3 sequences, at position 235 (C to T) and 322 (G to T) shows an important phenomenon which might be adopted by the virus to survive. As severe dengue is linked with the serotype-2, the genetic variations in this serotype points towards the much specific strategy to be adopted in near future to manage the severe dengue disease. In conclusion, we can say that, genetic diversity in the CprM region is present in the different serotypes circulating in the patients from Kerala India and this information may help in the management of dengue viral disease.

## 1. Introduction

Dengue viral disease is a constantly increasing mosquito transmitted infection which affect the populations globally [1; 2]. The incidences of dengue viral have increased about 30 folds during the last 50 years [3; 4; 5; 6]. The region for increased incidences of this disease are due to globalization, microevolution of the virus’s population, non-management of urban developments and patterns of climatic change [3; 7]. On an estimate, roughly half of the world population is at the risk of this dengue viral fever (WHO, 2009) [7; 8]. It is interesting to note that, the dengue reported cases is being doubled each year, and about twenty-five thousand people die annually due to this disease [9; 10].

While the disease pathogenesis of this disease is not known, no proper drugs or vaccine is available for the severe cases of the dengue fever [11; 12]. Dengue virus RNA is a positive-sense single-stranded and the virus is injected to humans by bites of mosquitoes i.e. *Aedes albopictus* and *Aedes aegypti* [13]. It is well known that dengue is caused by four different serotypes of dengue virus which are named DENV-1 to DENV-4, which are further divided into 19 distinct genotypes having various terminology among the researchers including the sylvatic types [14]. The first case of dengue was reported in Calcutta in 1963, since then many cases have been reported in the country with many out brakes through the country [14; 15]. There have been many cyclical trends of dengue outbreaks in the country, and now it is believed that this may attain an endemic disease status in India [14]. Kerala state is one of the predominant states in India having many numbers of out brakes of dengue and need intensive investigation. The problem of dengue viral disease particularly in Kerala is increasing since 2006 [15; 16]. It is important to note that, the Kerala state which is having ~1.5% land of the country, possesses 9.2% of dengue cases in 2010.

Viral mutation is a gradual and continuous process which happens in response to several pressures over a period [17; 18]. Therefore, a mutation specific study in the genome of the virus specially in the conserved region must be conducted to identity and estimate of the mutational process of the virus and their adaptability to caused disease in subsequent out brakes [19; 20]. Interhost dengue virus (DENV) have been considered to influence viral fitness and disease pathogenesis [21; 22]. Generally, few viral genes of DENV virus are used to understand the genetic variations [23; 24; 25]. it is important to note that the intrahost diversity was also observed in the antigenically different domains of virus [1; 2]. The genetic diversity, of the virus is due to the mutation in the virus genome[1; 2]. In this process of mutation virus species evolve their self for a better adapted survival and more strong pathogenesis [2; 26]. Our data will be elucidating the variations in CprM region of gene of various serotypes of dengue virus which may help in understanding the intrahost diversity and may play and important information in understanding the relationship between epidemiological and pathophysiological features of the disease [12; 27].

The epidemiology of various serotype stains of dengue virus in the many part of the India has been constantly changing [28; 29]. Due to ever increasing numbers of dengue cases it becomes vital to understand the genetic various in this virus circulating in a region at a particular time [28; 29]. Notably, there are is very less reports on the genetic diversity patterns of dengue virus from the Kannur district of Kerala in the year of 2019 due to various regions. Furthermore, a deep understanding about the molecular basis of genetic diversity of dengue virus in this region almost very limited. The genotyping of various serotypes of DENV, based on *CprM* has been widely used to understand and establish the molecular diversity of this virus [30]. The *CprM* gene based phylogenetic trees provide a similar topology and no discrepancy among the genotypes/sub lineages [30]. We adopted a CprM gene sequencing approach to identify the genetic mutations in this region [3; 23]. Using this approach, we sequenced CprM region of three different dengue serotypes and their mutational studied was defined [16]. Therefore, considering all the above facts mentioned above the present study was undertaken during August 2019-October 2019 to understand the genetic nature of circulating DENV in and around the Pariyaram medical college, Kannur, Kerala, India by sequencing the *CprM* gene of various serotypes [29; 30].

## 2. Materials and methods

### 2.1. Details of patients’ samples used in the study

Dengue fever suspected individuals visiting the **Pariyaram Medical College, Kerala India, hospital** which is recently named as **Government Medical College Kannur, Pariyaram,** OPD from Aug 2019 to October 2019 with an acute undifferentiated fever within 3 days of onset were tested using Dengue NS1 antigen rapid test. All the dengue NS1 positive individuals, were included in the study for the further confirmation of dengue by PCR and determination of serotype of dengue virus. For this purpose, 2 ml of venous blood samples was collected in EDTA vial and 1ml of blood in non EDTA vial and transported to ICMR Vector Control Research Centre, Puducherry by maintaining cold chain. Information on clinical symptoms and no of days from the onset of fever were recorded. The results of rapid antigen test were communicated to the individuals immediately and managed as per the hospital guidelines.

### 2.2 Ethical Approval

This study was approved by Institutional Human Ethics Committee (IHEC-0419/M) of ICMR- Vector Control Research Centre’s. The informed consent was obtained from all the study participants in writing. All the data obtained were kept confidential and anonymised.

### 2.3. Dengue virus detection by RT-PCR

Viral RNA was extracted from serum samples and using QIAmp viral RNA extraction kit (Qiagen, Germany) following manufacturer’s instructions. The RNA eluted was stored at 80 °C until further use. Titan One Step RT-PCR kit (Roche Diagnostics) was used to check the actual infection status of dengue virus using the primers targeting the capsid pre-membrane membrane region (Lanciotti et al., 1992). In this reaction viral RNA is reverse transcribed to cDNA and then to DNA in single tube using the enzyme mix which contains the reverse transcriptase and taq polymerase enzymes. The PCR reaction contains 5 μl of extracted RNA, 20 pM of each of the primers, and 5 μl of buffer, 0.5 μl of enzyme mix (Roche Diagnostics) and the reaction volume was adjusted to 25 μl. Reverse transcription PCR was performed at 50 °C for 30 min followed by an initial denaturation for 5 min at 94^0^C and 40 cycles of denaturation at 94^0^ C for 1 min, annealing at 50^0^ C for 1 min and elongation at 68^0^C for 1 min with a final elongation at 68^0^C for 8 min. The PCR products were run on agarose gel electrophoresis to verify the amplicon size of 511 bp

### 2.4. Serotyping of Dengue virus

The samples which were positive for capsid precursor membrane region of 511 bo were further subjected to semi–nested PCR to determine the dengue serotypes using the four dengue serotype primers TS1, TS2, TS3 and TS4 targeting the CprM region. The PCR reaction was set up as per Lanciotti et al., 1992 with the PCR conditions of 94^0^ C for 2.0 min followed by 25 cycles of denaturation at 94^0^ C for 1 min, annealing at 55^0^ for 1 min and extension t 72 for 2 min. The PCR products were electrophoresed in 2% agarose gel and was documented in GELSTAN gel documentation system.

### 2.5 Sequencing of dengue virus serotypes

The serotyped PCR products visualised under UV light were cut from the agarose gel using x-tracta Tool (Sigma Aldrich). The excised DNA fragments were further refined using nucleospin gel and by using PCR clean-up kit (Macherey-Nagel) and quantified for determining the purity and concentration. Cycle sequencing of the different serotyped DNA bands was accomplished using BigDye™ Terminator v3.1 Kit used for Cycle Sequencing (Thermo fisher Scientific). The cycle sequenced products were purified by NucleoSEQ columns (MAcherey-Nagel) and analysed in Genetic analyser 3130Xl (Applied Bio systems)

### 2.6. Sequence analysis

Sequences obtained were edited using Chromas pro V.2.1.8 and blast search against the GENBANK database. The blast hit sequences were aligned using Bioedit V 7.0.5.3 to check for the variation in the sequences. Phylogenetic analysis of the different serotype sequences obtained from this study was performed along with the genbank dengue serotype isolates using Mega V.7.0. Dengue virus sequences obtained from Dengue positive samples in the present study were blast searched and the blast hit sequences retrieved from GenBank databases were subjected to multiple sequences alignment using Muscle. Phylogenetic tree was constructed using Maximum Likelihood method based on Tamura Nei model in MEGA 7.0 software. The robustness of the tree was assessed with 1000 bootstrap replicates.

### 2.7. Statistical analysis

The data is represented as per the requirements of the experiment or otherwise specified. The software GraphPad prism 5 was taken in use for analysing data wherever needed.

## 3. Results

### 3.1. Dengue Virus detection and serotype determination

Of the 25 samples positive for dengue NS1 antigen, samples were positive for dengue virus by RT-PCR assay. All the samples positive by reverse transcriptase assay were serotyped and of the samples. All the four serotypes were found in the samples cDNA **synthesis and synthesis and multiplexing of PCR gives four different serotypes of DENGUE virus.** Dengue NS1 positive samples were subjected to Reverse Transcriptase PCR targeting Capsid Precursor membrane (CprM) region using the primers specific to this region. An amplicon size of 511 bp was amplified. Dengue serotyping was performed using the primers specific to CprM region which can distinguish all the four serotypes of dengue by a Semi nested PCR. Of the 25 samples 18 samples were found confirmed with dengue infections (Table No. 2). The four distinct serotypes were present in the samples (Fig. No. 1). Serotype specific amplicons were excised from the agarose gel and cycle sequencing reaction was accomplished by Big dye terminator cycle sequencing kit, V.3.1. The sequences were edited using Chromas Pro. Forward and reverse sequences were assembles using Bio Edit software and the sequences were blast searched against the Genbank nucleotide database. DENV-1, DENV-2 and DENV-3 were found to be 100% identical to the respective serotypes in the GenBank nucleotide database while DENV-4 alone had variation in the sequences. This variation needs to be confirmed by multiples sequence alignment with other DENV-4 sequences. The accession numbers of GenBank are OM510975, OM510976, OM510977, OM510978, OM542335, OM572553, OM572554. These are CprM gene sequences obtained in the present study.

**Figure No.1.**
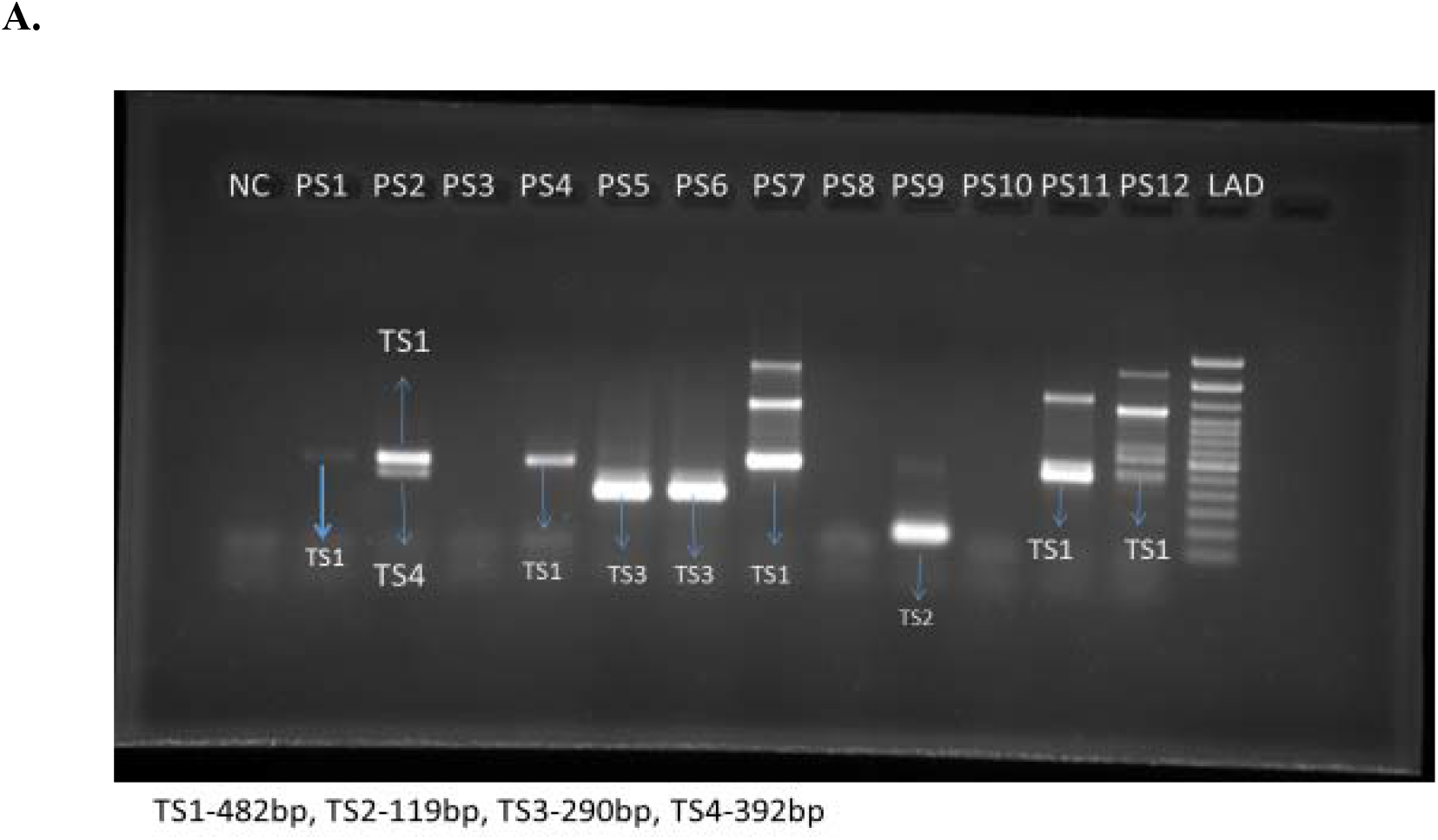
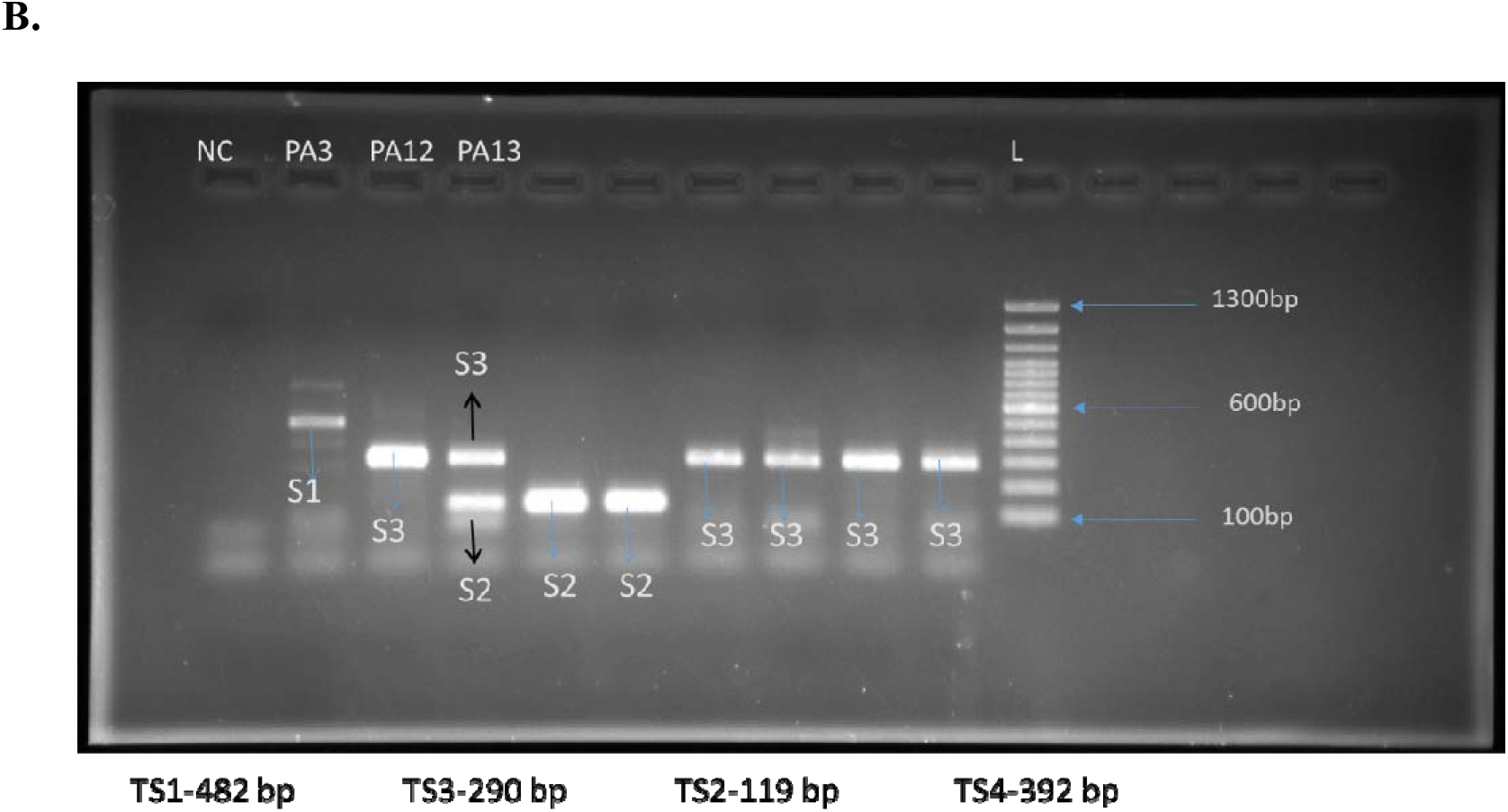
Determination of specific serotype present in the samples by semi nested PCR technique which is followed by the agarose gel electrophoresis for the identification of proper size of amplicon. as shown in the figures A, B, C, serotype-1 (TS1) 482bp, serotype-2 (TS2) −119bp, serotype-3 (TS3) −290bp, serotype-4 (TS4)-392 bp. **A.** Determination of different serotypes of dengue following the protocol of Lanciotti et al. 1992, with desired modifications. As described in the figure we have got all four serotypes of dengue. the percent calculation of dengue is also derived and presented elsewhere. Figure also showing one of the representative co-infections of serotype 1 (ST1) with serotype-4, ST-4. 100 bp ladder was used to determine the size of the amplicon. **B.** agarose gel image showing one of the coinfections present in the samples of serotype 2 (ST2) with serotype-3, ST-3.

### 3.2. Assessment of percent prevalence and coinfection among the positive samples obtained by the serotyping

Before analysing the serotypes, we have done diagnostic PCR and confirmed that, which samples is positive for dengue or not. Out of total samples obtained from the Kannur medical college which were confirmed, NS1 positive, rapid antigen test, only 77% were found positive for dengue using PCR. All the four serotypes were found to be present in the sample. Serotype 1 and 3 was the predominant type (33.3 %) compared to other serotypes. As we know that co-infection of dengue virus is an important phenomenon to describe the intrahost compatibility and fitness of different serotypes of dengue virus. Further, to understand the coinfection percentage prevalence (Fig. No. 2 A and B), we have analysed our serotype specific data and found that a small number of serotypes also present among the positive samples. Coinfection of serotype 1 and 4 was highest percentage (11.1 %) compared to serotype 2 and serotype 3 (5.5 %).

**Figure No.2.**
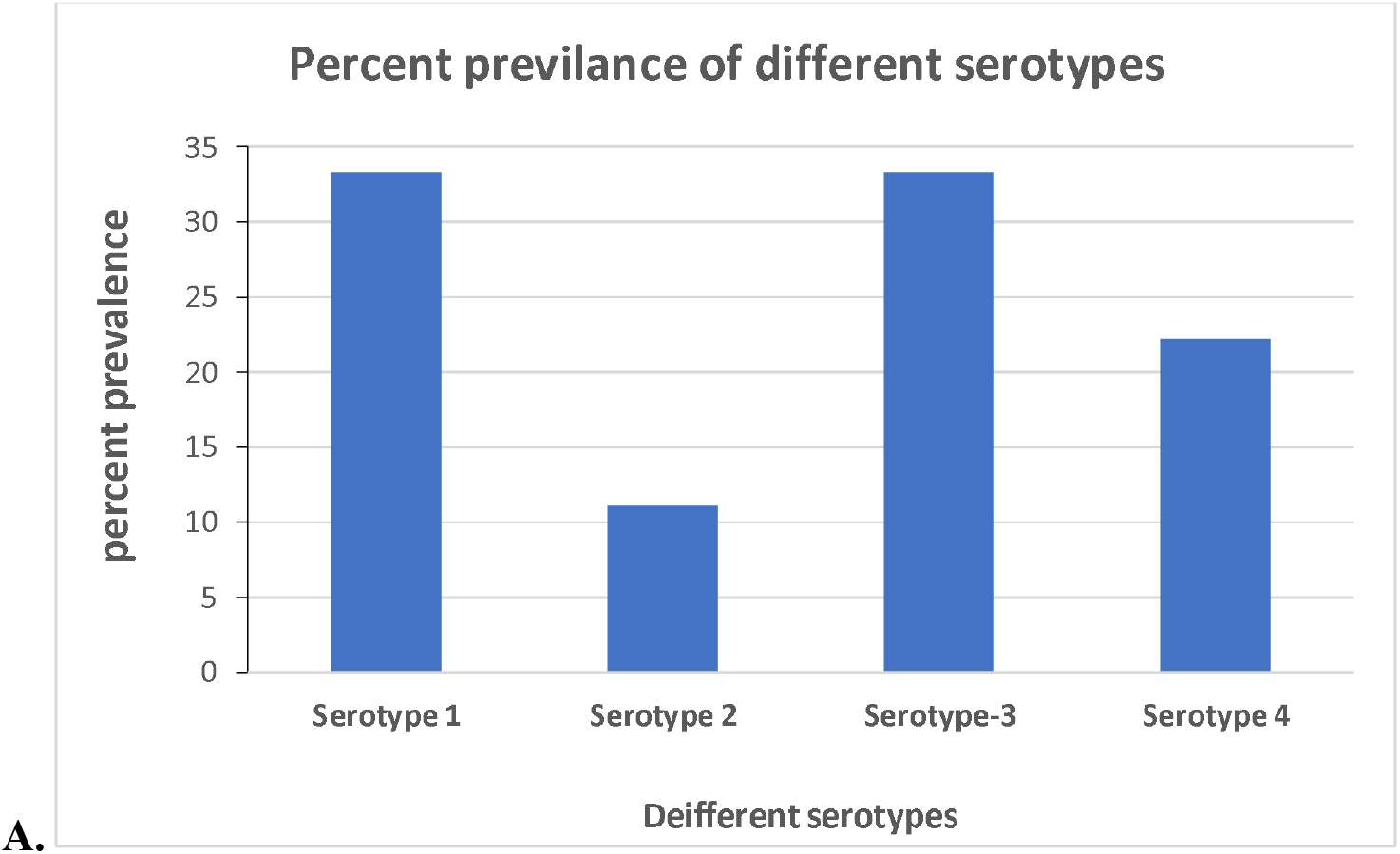
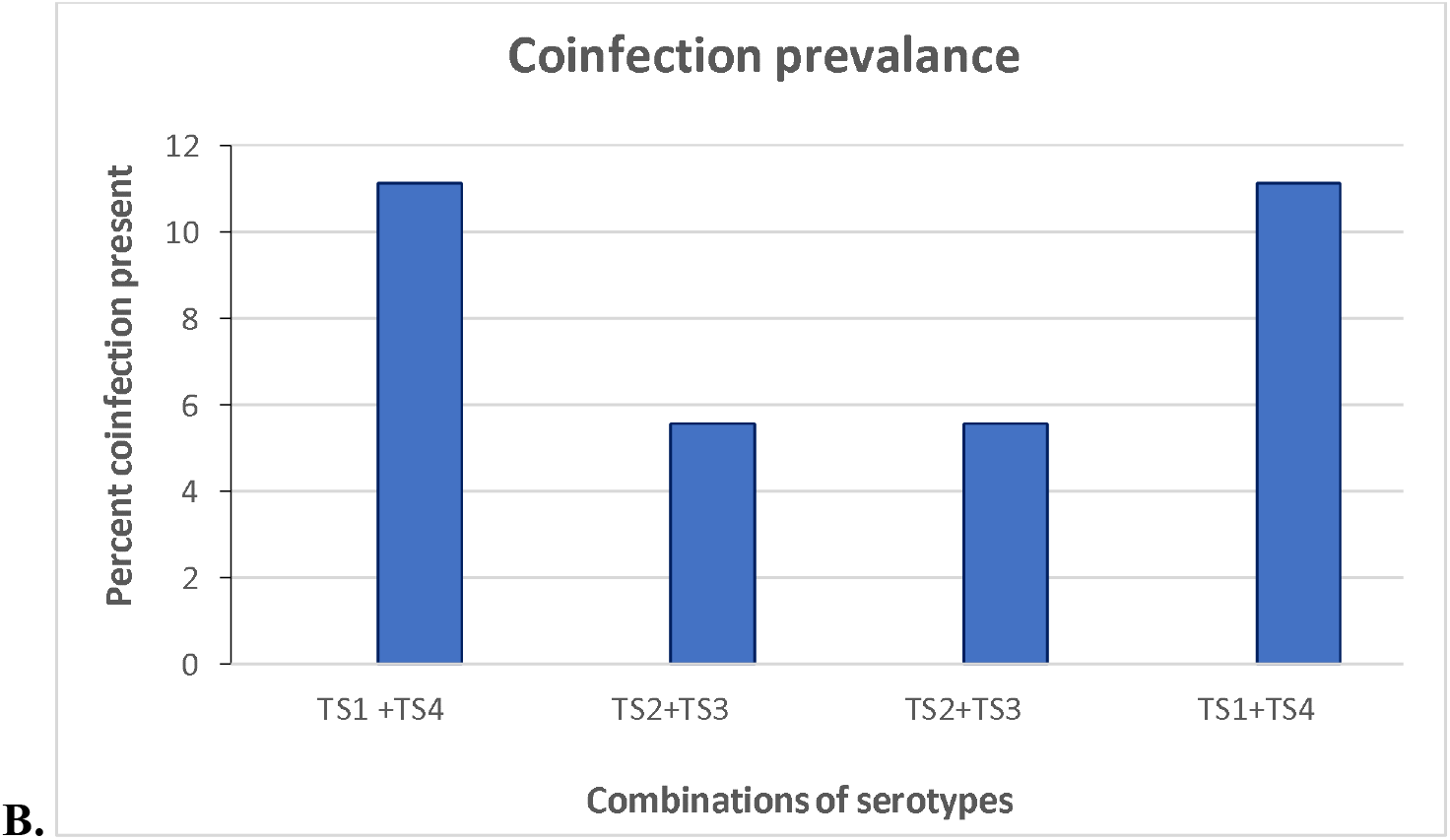
**A.** Percent prevalence of different serotypes present in the patients coming to the Pariyaram medical college Kerala during the august-October 2019. B. Percent prevalence of co-infections of serotypes present in the patients coming to the Pariyaram medical college, Kerala during the august-October 2019.

### 3.3 Assessment of genetic mutation in the CprM region and their correlation

Genetic mutations in virus genes are very important which are very important for virus to adapt in various situations [31; 32; 33]. Therefore, next, in order to categorise the phylogenetic similarity and dissimilarity we have constructed a phylogenetic tree and placed our sequences [34]. Phylogenetic tree constructed with dengue virus sequences obtained from dengue virus positive patients showed greater than 98.97 % similarity with GenBank isolates and the DENV-4, DENV-3 and DENV-2 sequences formed separate cluster. Two different strains were found among the DENV-3 sequences and were found to cluster separately with GenBank isolates. This is due to the variation in the nucleotide sequence of Cap PrM gene. Sequence variation among the four DENV-3 sequences were shown at position 235 (C->T) and 322 (G->T) and revealed that 2 of sequences were like Delhi and Pune isolates while the other 2 were like Mumbai isolates (Fig. No. 3). These variations did not result in amino acid change and of synonymous mutations.

**Figure No.3.**
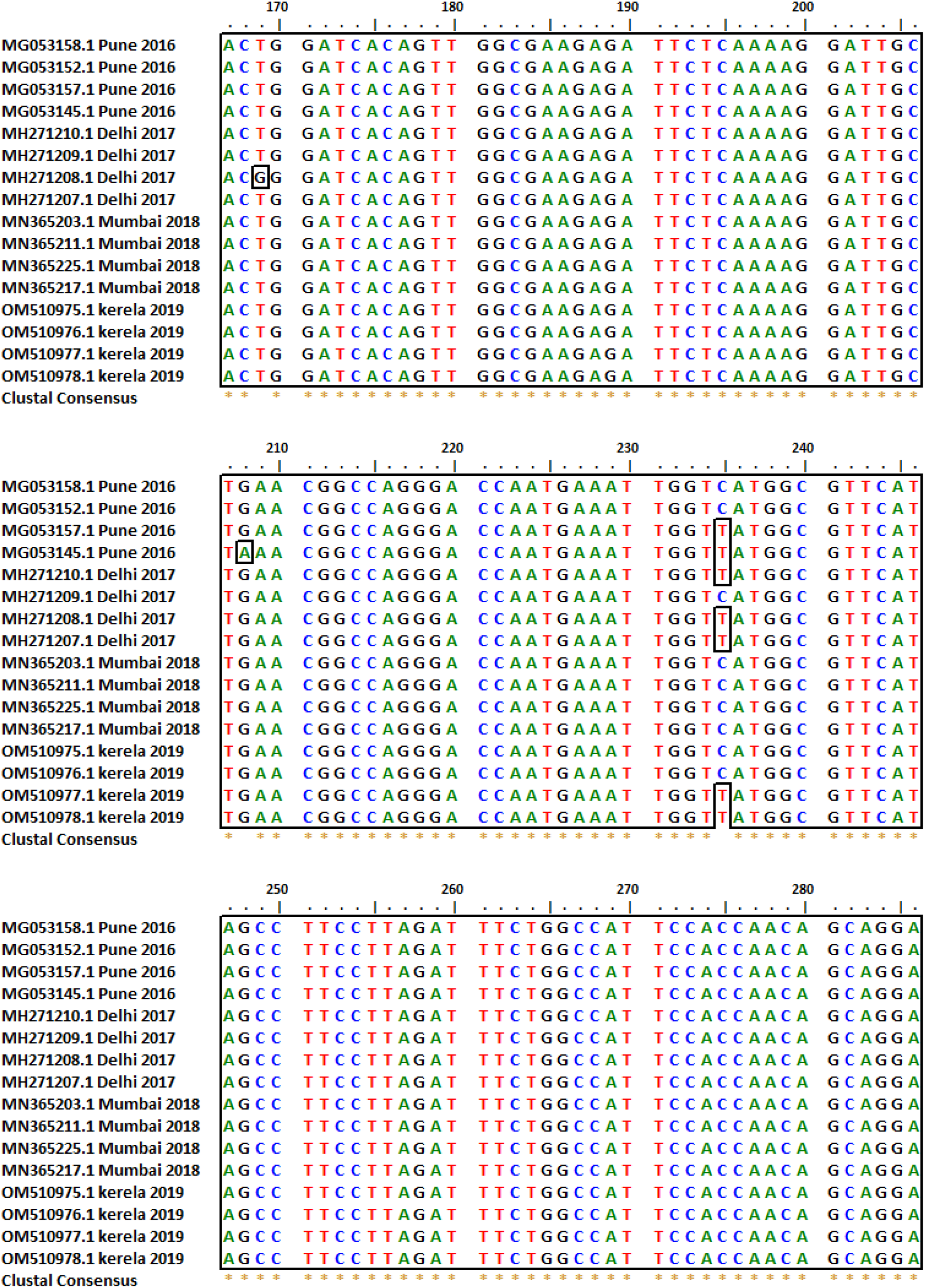
**Fig.** Sequence alignment of the CprM gene sequences of DENV-3 isolates obtained from the study (OM510975.1 to OM510978.1) with GenBank DENV-3 isolates of India circulating during 2016-2018. Sequence variation among the four DENV-3 sequences were shown at position 235 (C->T) and 322 (G->T) and revealed that 2 of sequences were similar to Delhi and Pune isolates while the other 2 were similar to Mumbai isolates.

### 3.4. Analysis of gene mutations to their amino acid change in the protein and their anti-genic analysis

Virus fitness also depends on the structure of the virus protein capsid and outer membrane which in turns depends on the amino acid change in the virus protein [35]. We have further analysed the all serotypes mentioned that is there is any change in the amino acid sequence of the serotype or not [35]. We found that, there is almost no change in the amino acid change in the all three serotypes of the virus. Phylogenetic analysis of the obtained serotypes and their similarity with existing serotypes. In addition to the protein mutation we have also assessed the phylogenetic analysis of the sequences we obtained that, these serotypes are somewhat closely related to the existing reported genotypes in India and no new genotypes was observed [34] (Fig. No.).

### 3.5. Phylogenetic analysis of all the serotypes obtained in the study

After the identification of the serotypes, we have done the genotypic analysis of each serotype (Fig. No. 5 to Fig. No. 8). In the molecular phylogenetic analysis, by maximum likelihood methods the bootstrap support value suggested that dengue serotype 2 (Gene bank accession no. OM572553, isolate DENV-2_P9, and OM572554_ isolate DENV-2_PA15) shares a close relationship with the Genotype-4. It is important to note that, DENV-2 was previously closer to a different clade which has been now shifted. The serotype-1 fall in the genotype-5 which is also showing some little difference from the existing strains reported from the various regions from India. Similarly, Dengue serotype-3 isolates (gene bank accession numbers, OM510975_ isolate DENV-3_PA8, OM510976_ isolate DENV-3_PA12, OM510977, isolate DENV-3_PA19 and OM510978_ isolate DENV-3_PA21) fall under the genotype-3 and a little genotypic shift is seen in this type 3 serotype of dengue virus. The type 4 serotypes with the (GenBank accession no. OM542335_ isolate DENV-4_P2) falls like the genotype-I but having distinct culture and also little deviate from existing reported dengue viruses type −4.

**Figure 4.**
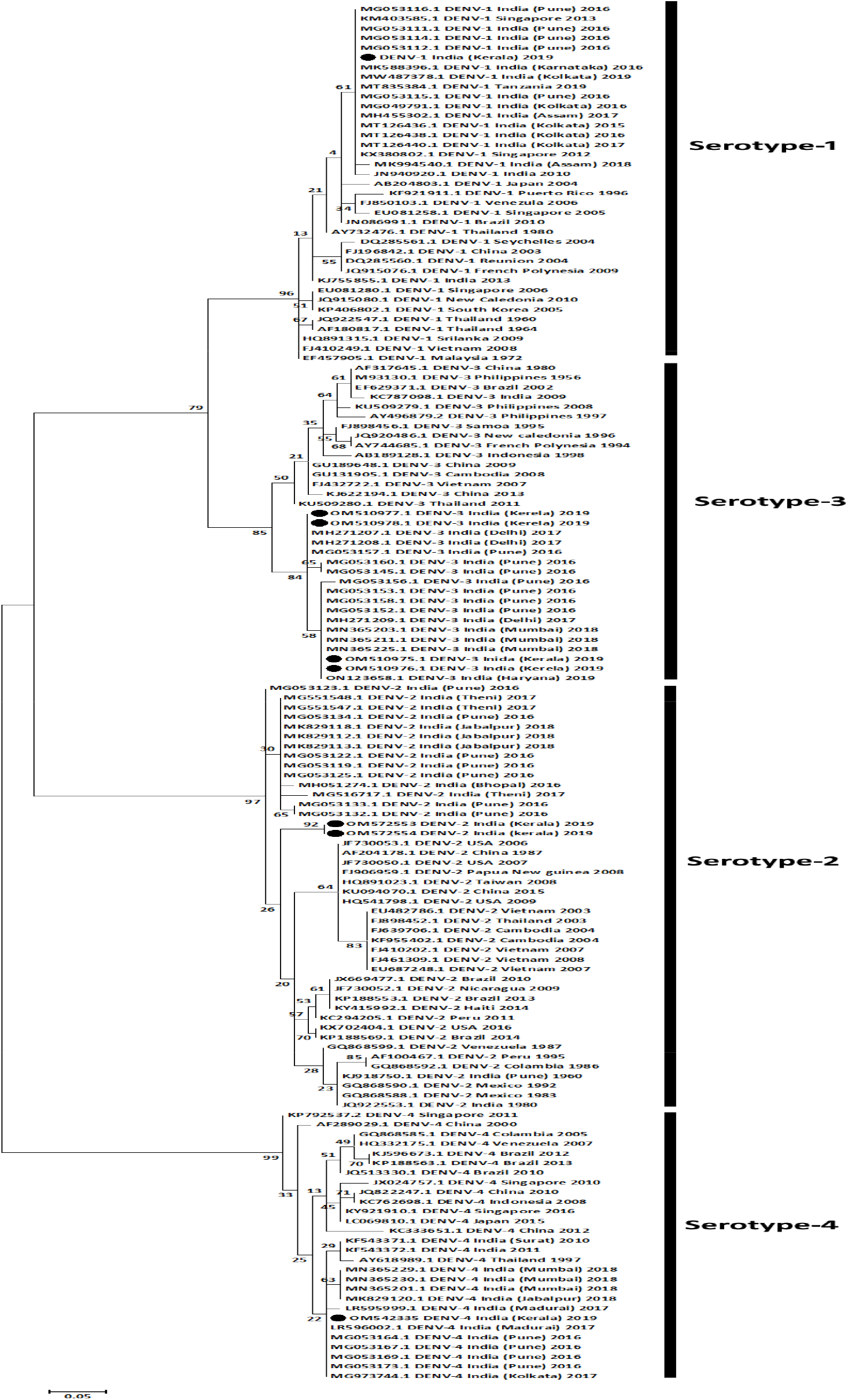
Phylogenetic analysis for the determination of different **serotypes** using CprM gene sequences of all the four dengue serotypes obtained from this study and other DENV sequences from genbank database. Genbank sequences are indicated by the Genbank accession number, country and the year of isolation. Sequences from this study are indicated by black circles. Scale bar indicates the number of nucleotide substitutions per site.

**Figure 5.**
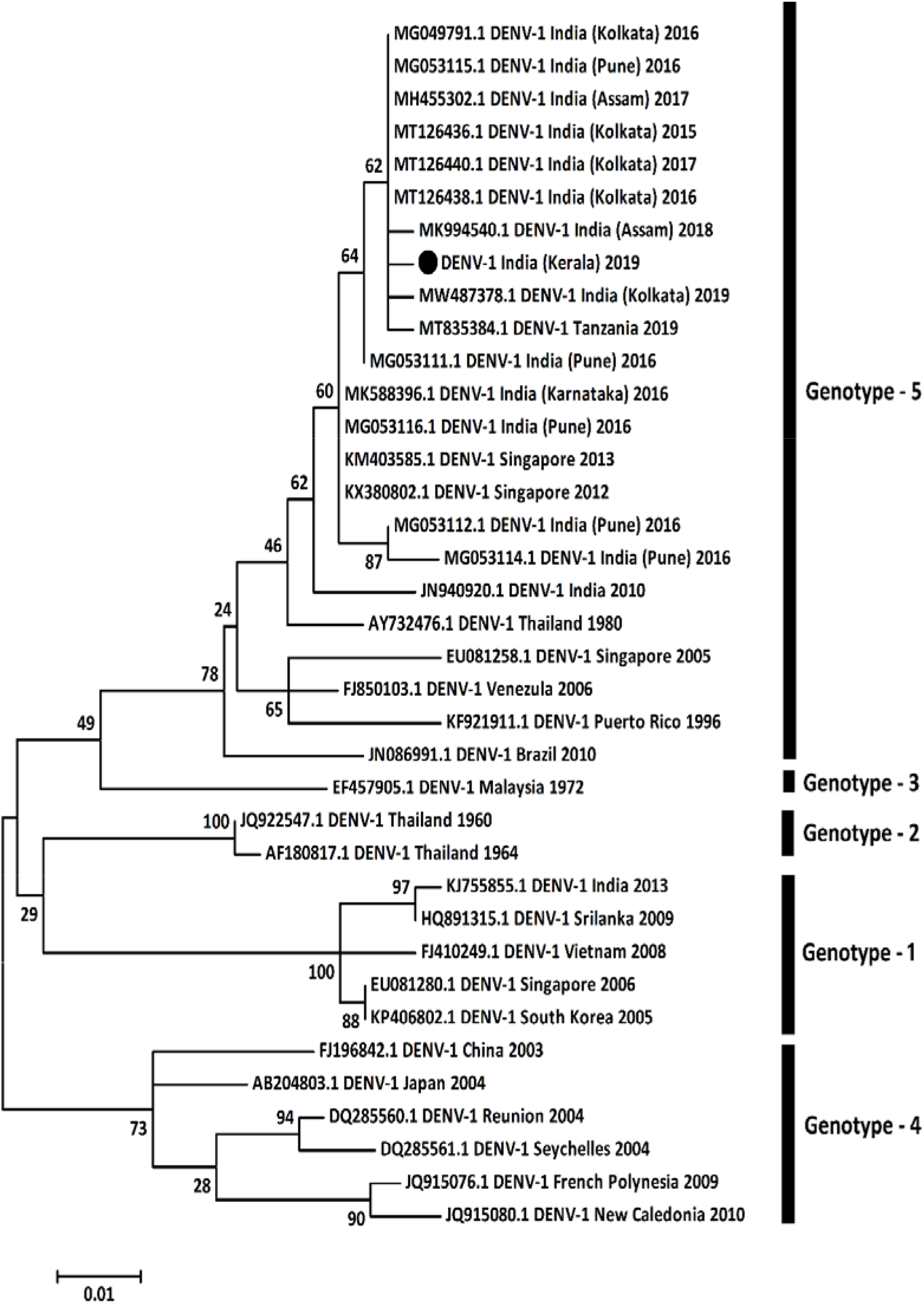
Phylogenetic analysis for **genotypic** determination of **DENV-1 serotypes** using CprM gene sequences of obtained from this study and other Genbank sequences (submitted latter). Genbank sequences are indicated by the Genbank accession number, country and the year of isolation. Sequences from this study are indicated by black circles. Scale bar indicates the number of nucleotide substitutions per site.

**Figure 6.**
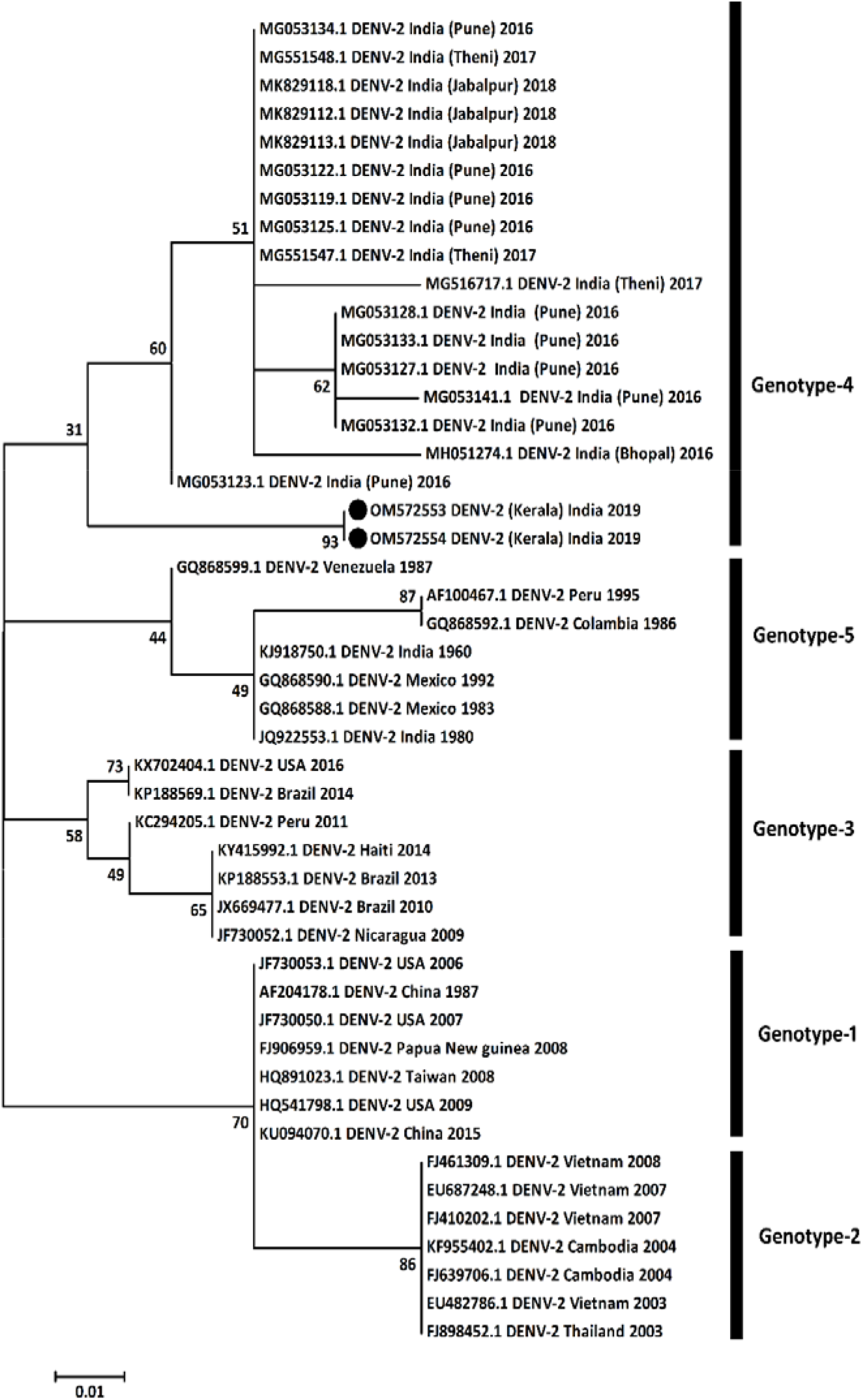
**The genotypic determination** of **DENV-2 serotypes** using CprM gene sequences obtained from this study and other Genbank sequences. Genbank sequences are indicated by the Genbank accession number, country and the year of isolation. Sequences from this study are indicated by black circles. Scale bar indicates the number of nucleotide substitutions per site.

**Figure 7.**
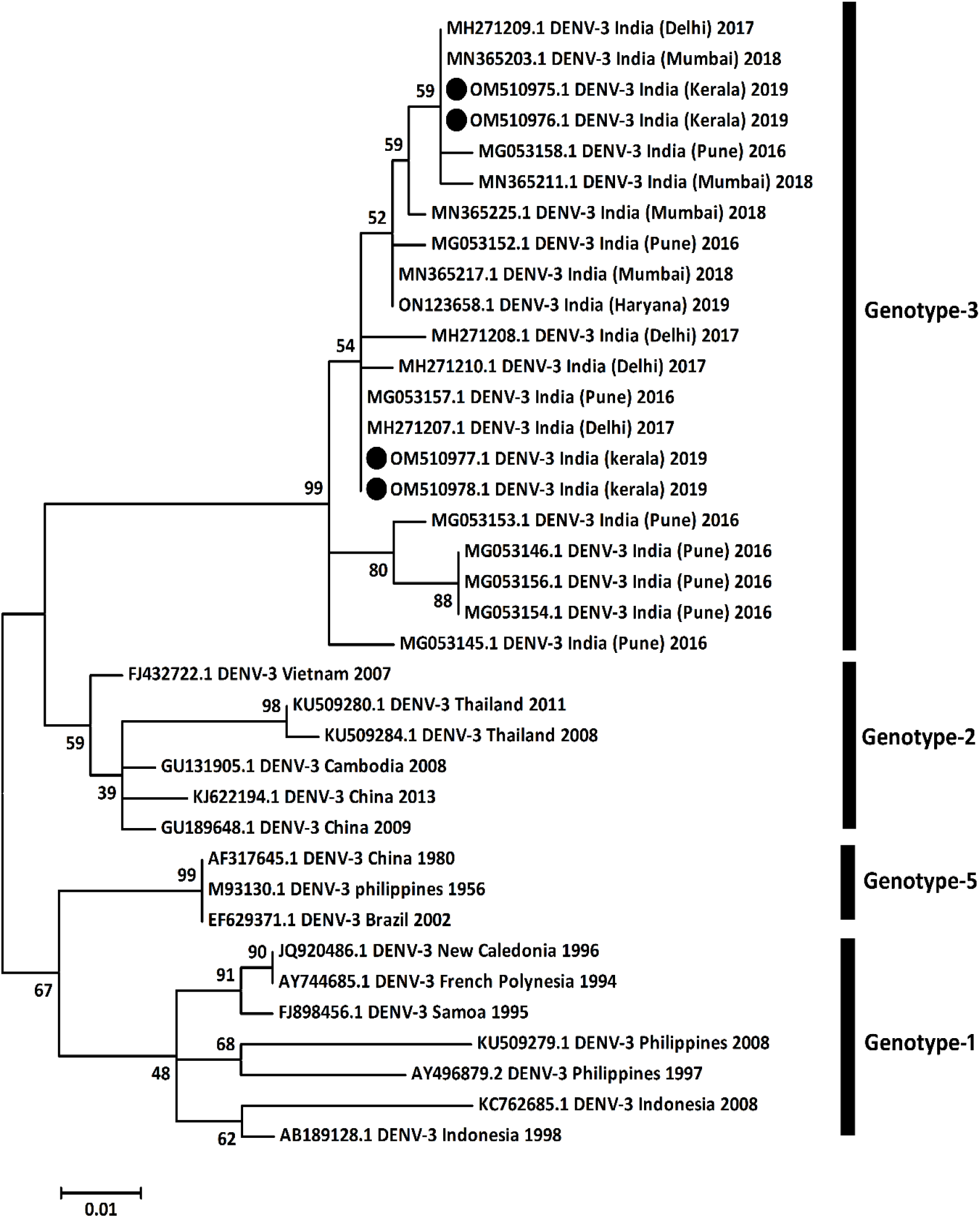
Phylogenetic analysis for the determination of genotypes using CprM gene sequences of **DENV-3 serotypes** obtained from this study and other Genbank sequences. Genbank sequences are indicated by the Genbank accession number, country and the year of isolation. Sequences from this study are indicated by black circles. Scale bar indicates the number of nucleotide substitutions per site.

**Figure 8.**
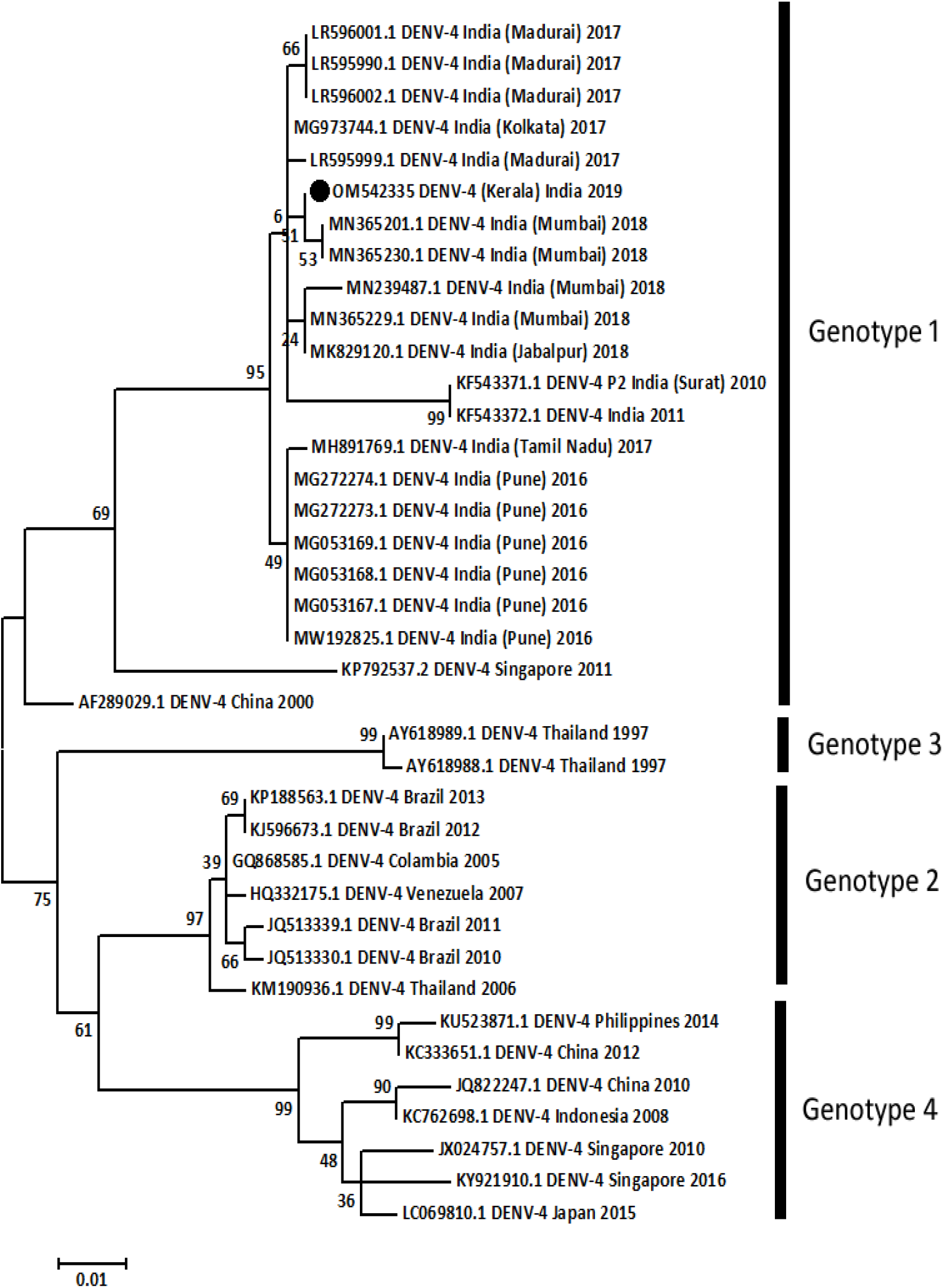
Phylogenetic analysis for the determination of genotypes using CprM gene sequences of **DENV-4 serotypes** obtained from this study and other Genbank sequences. Genbank sequences are indicated by the Genbank accession number, country and the year of isolation. Sequences from this study are indicated by black circles. Scale bar indicates the number of nucleotide substitutions per site.

## 4. Discussion

In the present study, we have accomplished CprM region sequencing of the dengue virus to understand the prevalence and genetic diversity of the dengue virus in the patients from Kerala, India [27; 36]. This CprM gene-based sequencing may clarify the various genotypes and serotypes of dengue virus prevalent in a region at a time [15; 37]. The information generated in this study helped understand the gene-specific differences of dengue virus among the different patients to elucidate the linkages with disease severity and the pathogenesis [38; 39]. It is important to note that, Dengue virus (DENV) is an arbovirus and transit from human to mosquito and vice versa [39]. How, DENV adapt in the different hosts is an important question and the answer perhaps lies in the gene mutations. Therefore, we used CprM region sequencing and estimated single-nucleotide variants in these virus strains of different serotypes [40; 41]. This allowed us to determine the mutations to the collective fitness of the viruses [39].

There have been many out brakes in Kerala of dengue viral disease with a major one in the year 2003. Since then all districts of Kerala observe many out brakes during the various rainy seasons. It is surprising to see, since the year 2007 there have been many seasons in which the dengue cases are continuously increasing with a maximum increase in the year of 2010. Notably the Kerala state has now become the thirst most affected state of the India. The *Aedes albopictus* mosquitoes are a plenty population in the state of Kerala. In this study, we obtained the **four distinct serotypes** prevalent in and around the Pariyaram medical collage Kannur, Kerala State. We found the DENV-1 and DENV-3, serotype circulating in Kerala is considered endemic in Kerala and with little to no information on its status in Pariyaram, Kerala India [42; 43]. In every season the main source of information of different serotypes are from field caught mosquitos and serological surveys in humans.

It is very important to note that, currently only licensed vaccine, CYD-TDV (known by trade name Dengvaxia and made by Sanofi Pasteur, Lyon, France) needs serostatus of the area and prevailing dengue serotypes [44]. This makes the importance of existing serotypes in the area as safety issues linked with higher rate of hospitalization may happen for individuals who have never been infected with dengue before [45]. Further in is important to note that, coinfection with more than one serotypes during dengue viral disease enhances the complexity for the diagnosis of the virus. Another important mechanism of the dengue pathogenesis is antibody dependent enhancement or ADE [45]. The ADE occurs when patients contract a heterotypic secondary dengue virus infection and make additional complication for effective treatment of patients [46]. The development of Directly Active Antivirals (DAA) are an important area where scientists are trying to make some anti-dengue drugs which can suppress the all kinds of genetically diverse genotypes and/or serotypes of dengue virus [47; 48]. However, tremendous efforts of the scientists to find out some antiviral therapy, there is no anti dengue drugs in the market presently. It is well known that; beneficial mutations generally map to intrinsically disordered domains of the virus which are clustered to a specific part in the genome [2; 49]. Several studies demonstrated that, the continuous influx of several mutations in the direction of virus adaptation [22; 40]. Further, it is important to note that, phenotypically redundant adaptive alleles together enhance the host-specific DENV adaptation [50].

It has now been widely adapted that coinfection with dengue serotype alone or in combination with different virus cause immunocompromise stage and or the immune dysfunctions resulting in the more severe complications [51; 52]. In the present manuscript we have obtained the co-infections which in turn may points towards the heterogeneous ADE in dengue infections [53]. In the present, study we have obtained that, coinfection of serotype 1 and 4 was highest percentage (11.1 %) compared to serotype 2 and serotype 3 (5.5 %) [54]. This co-infection and its combination is still less studied in terms of immunopathogenesis and need extensive research is needed to further clarify the findings, however, some of the studies have corelated it with the severity of the disease [55]. It is important to know that the similar to the present study the coinfection (serotype-3 with serotype-2) was observed in the Cameron in 2021 which was suspected towards increased symptoms and severity of disease [54; 56]. In another study from the brazil, it was reported that, serotype-1 is more able to case severity compared to other serotypes, suggesting an important aspect of comparative severity among four distinct serotypes [57]. The immune dysfunctions and immune compromise stages in coinfections are considered due to various reasons. One such region towards immunocompromise stage is that, it impairs the neutralizing antibody and CD4^+^ T Cell responses which indirectly allows the one virus over the others. The different time of infections and antibody dependent enhancements (ADE) are also considered to be an important aspect in this concept of disease severity and immunopathogenesis.

As explained in many studies, Dengue virus mutation is an important phenomenon in terms of virus survival and to cause the disease severity [58]. In the present study, it was obtained in DENV-3 sequences that, mutation at position 235 (C to T) and 322 (G to T) shows an important phenomenon which might be adopted by the virus to survive in the adverse conditions for under the environmental pressure [58]. This mutation is not causing any change in the protein amino acid and thus called synonymous [59]. In many study of viruses it has been found that, even the synonymous mutation can change the virus adaptability and antigenicity, however the exact mechanism and understanding is still debated in this aspects and need extensive investigations [60]. Another aspect of synonymous mutation is the choice of codon by the dengue virus among its natural hosts is also described in many studies. It is described that, the dengue virus has the evolutionary distance among its natural hosts (*Homo sapiens, Pan troglodytes*, *Aedes albopictus* and *Aedes aegypti*) [61]. In further studies, it needs to be further analysed that these synonymous mutations are of which purpose and may be used as a source of information.

## Conclusion

The present study describes the prevalence of all the serotypes (DENV1 to DENV-4) circulating in patients in and around the Pariyaram medical college, Kerala, India. It was important to note that, a high level of mutations was observed in the CprM gene giving rise a high level of genetic diversity in all 4 different serotypes of dengue virus in this area. This suggests that a continuous monitoring of the dengue virus serotype is needed in this region for better prevention and therapeutic strategies. The genetic mutation also provides me with the phylogenetic understanding of various genotypes present in a time in this area of Kerala India. We also report that, co-circulation or co-infection of all four serotypes of DENV in Pariyaram district of Kerala, in the combination of DENV1 along with DENV-3 and DENV-2 in the combination with DENV-4. The results in this article provide a clear-cut understanding of dengue virus diversity in Kerala state of the India, however further study in this area would be vital and provide more deeper understanding of the dengue viral disease.

**Table No. 1.**
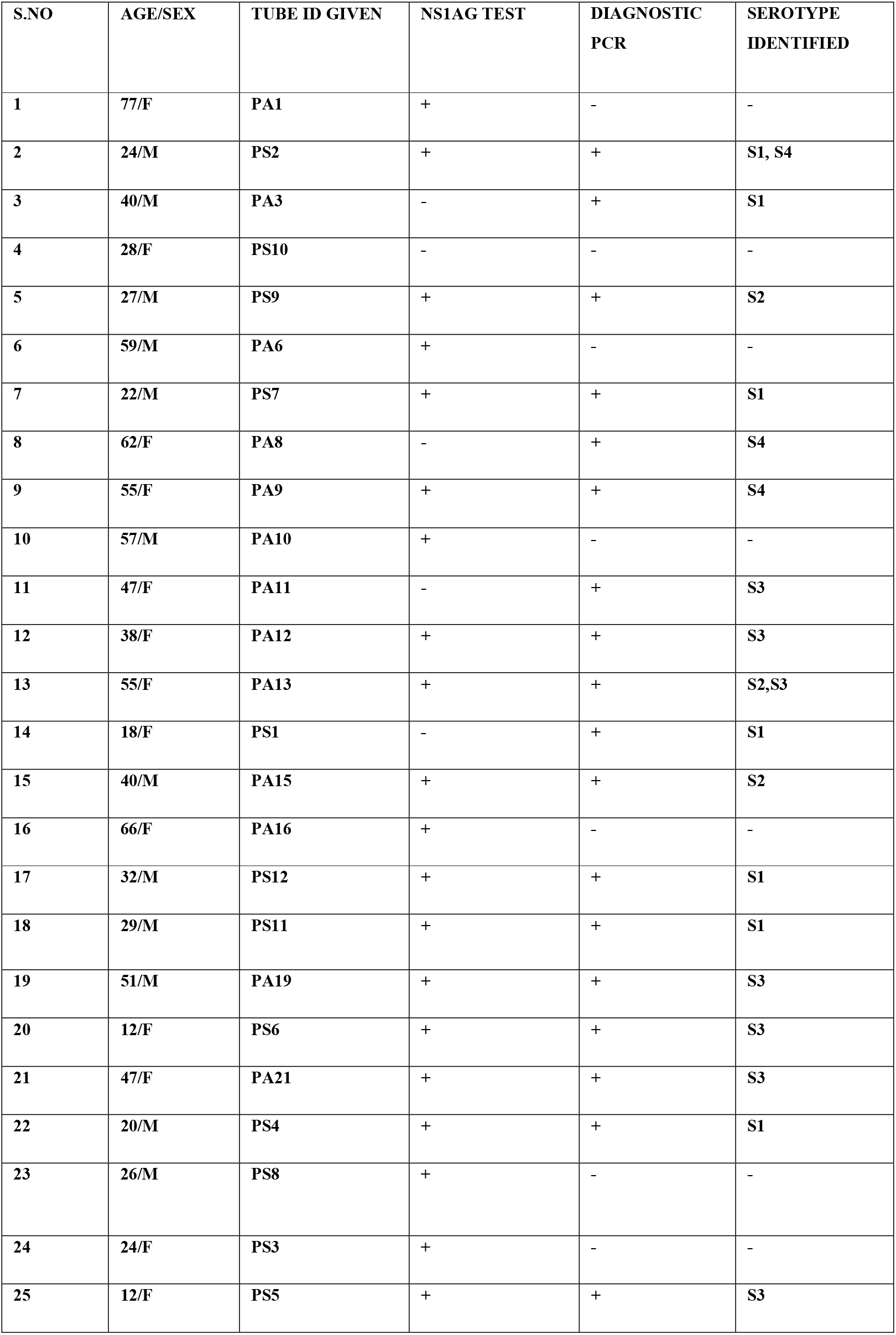
Details of the patient’s samples along with their respective serotypes.

## Abbreviations

NS: non-structural protein
CprM: Capsid precursor Membrane
DENV: Dengue virus
TS1: Type-1 serotype
TS2: Type-2 serotype
TS3: Type-3 serotype
TS4: Type-4 serotype
DHF: Dengue Haemorrhagic Fever
DSS: Dengue Shock Syndrome
DCs: dendritic cells
MMPs: matrix metalloproteinases
CHIKV: chikungunya virus
CD4: cluster of differentiation 4
CD8: cluster of differentiation 8

## Acknowledgements

Grant No: 61/16/2020-IMM/BMS from Indian Council of Medical Research, New Delhi, India, to Dr. Rituraj Niranjan, entitled “Dengue Shock Syndrome (DSS): study on the role of blood matrix metalloproteinase-14 (MT1-MMP/MMP-14) associated to innate immune cells and its contribution to endothelial dysfunctions” as an extramural Ad-hoc grant is also gratefully acknowledged.

## Availability Data

The data presented in this manuscript will be available to others based on the valid request.

